# The role and effects of the phage T4 Ac protein on infection

**DOI:** 10.1101/2025.06.30.662393

**Authors:** Ana G. Arvizu Santamaría, Steven N. Rosvold, Fabini Orata, Anastasia L. Elias, Dominic Sauvageau

## Abstract

The *ac* gene from T4 and T4-like phages is associated with susceptibility to acriflavine, an acridine that intercalates with DNA, disrupting replication and transcription. While this interference has been shown to inhibit phage replication – resulting in reduced frequency of infection and viral titer at the population level – and mutation or knockout of the *ac* gene renders the phage resistant to acriflavine, the role of the *ac* gene product, Ac, has not yet been elucidated. This study aims to further explore the role of the *ac* gene in phage infection by expressing it in *Escherichia coli* and evaluating its effects on the host under varying growth conditions and during phage infection, both in the presence and absence of acriflavine. *E. coli* induced to express the *ac* gene from phage T4 showed an increased susceptibility to acriflavine compared to the same strain not undergoing expression (non-induced) or not carrying the *ac* expression plasmid (empty). Moreover, in the presence of acriflavine, the parental phage T4 was not able to infect the host variants (induced, non-induced and empty), suggesting the Ac protein is involved in a potential membrane modification leading to acriflavine hindering infection. When the *ac* gene was deleted from the T4 phage (T4Δ*ac*), the mutated phage was capable of infecting the three host variants (empty, non-induced and induced) in the presence of acriflavine, showing resistance to this acridine. Based on experimental results and protein structure prediction, we propose that the *Ac* protein integrates into the bacterial host cell membrane and interacts with the AcrAB-TolC efflux pump, either altering its conformation or blocking its function, thereby preventing the excretion of acriflavine which accumulates in the cell and impedes DNA replication

## Introduction

Bacteriophage (phage) T4, a member of the Caudovirales order, represents one of the most extensively studied and genetically characterized viruses in molecular biology. Discovered in the first half of the 20^th^ century, T4 has since become a pivotal model system for understanding the intricacies of viral replication, host specificity, and the molecular mechanisms underlying phage life cycles (Kutter and Sulakvelidze, 2004; Hendrix et al., 2013). Its complex structure, elaborate genomic architecture, and sophisticated infection strategies have made it an exemplary subject for exploring fundamental concepts in virology and genetics (Wood and Revel, 1976; Hendrix et al., 2013). This lytic phage employs a series of meticulously orchestrated steps to hijack the host’s cellular machinery, culminating in the lysis of the bacterial cell and the release of progeny virions (Novik et al., 2017; Ge et al., 2020). One of the most remarkable features of phage T4 is its genetic robustness and adaptability, underscored by its ability to carry out sophisticated mechanisms of gene regulation, DNA repair, and host interactions (Miller et al, 2003; Petrov et al., 2010).

Additionally, the robust DNA replication machinery of T4 and innovative strategies for overcoming bacterial defences offer invaluable insights into viral evolution and host-pathogen interactions (Labrie et al., 2010; Ge et al., 2020). However, one puzzling feature of phage T4, and many T4-like phages, lies in their *ac* gene, which is linked to reduced infection in the presence of acriflavine and other acridines —a group of synthetic dyes with antiseptic, antimicrobial, and anticancer properties (Foster, 1948; Silver, 1964; Schmidt and Liu, 2015). These compounds, which also include proflavine, intercalate into nucleic acids, disrupting DNA replication and transcription (Gatasheh et al., 2017). High-resolution structural studies have revealed how acriflavine binds to DNA, providing a clearer picture of its inhibitory effects (Manivannan et al. 2013; Mukherjee and Sasikala, 2013; Hasanzadeh and Shadjou, 2016). This mechanism is noteworthy when considering the effect of acriflavine on phages, which rely on host cellular machinery to replicate their genomes. For example, the fact that the intercalation of acriflavine into the host and/or viral DNA could potentially interfere with phage replication and assembly, decreasing the viral titer, was identified decades ago (Foster, 1948).

The susceptibility of T-even phages to acridines has been linked to specific genetic elements; the *ac* gene in T4 and related phages, and the previously named *pr* gene in T2 and T2-like phages (Hessler, 1963; Silver, 1964, 1967), now also referred to as *ac* gene in the NCBI database. The *ac* gene, located within one of three gene clusters encoding for predicted membrane proteins within the T4 phage genome, consists of 156 base pairs and encodes a protein of 5.5 kDa, composed of 51 amino acids. This gene is categorized as non-essential, yet its role in acriflavine sensitivity is notable (Silver, 1964; Miller et al., 2003). Despite all this information, the precise mechanism of action of the *ac* gene product, Ac, is not fully elucidated, although early studies have suggested to involve an early, energy-dependent modification of the bacterial outer membrane that increases acriflavine uptake (Silver, 1964, 1967). Naturally-occurring mutations in the *ac* gene leading to an unfunctional protein or complete removal of the gene (Δ*ac*) results in mutants that exhibit increased resistance to acriflavine (Hessler, 1963; Silver, 1964) Other mutations in T4 genes, such as *ama*—whose function remains undefined— and genes *60* and *39*—the two subunits of DNA topoisomerase—have also been associated with acriflavine resistance (Miller et al., 2003).

In many bacterial species, acridines are handled by efflux pumps (EPs), highly conserved integral membrane protein complexes that function as transporters, recognizing and removing toxic compounds from the inside of the cell to the outside environment (Amaral et al., 2014; Teelucksingh et al., 2020; Nishino et al., 2021) EPs provide resistance to multiple classes of antibiotics by transporting a wide range of substrates with different physicochemical properties, a phenomenon known as multidrug resistance. Members of the resistance-nodulation-cell division superfamily, which are commonly found in Gram-negative bacteria, play a significant role in clinically relevant antibiotic resistance. Notable examples of resistance-nodulation-cell division efflux systems include AcrAB-TolC in *Escherichia coli* and MexAB-OprM in *Pseudomonas aeruginosa*, which are well-studied and frequently associated with antibiotic efflux (Ma et al., 1994; Nikaido, 2011; Teelucksingh, Thompson and Cox, 2020) However, these EPs also recognize and expel various other compounds that are unrelated to antibiotics, including dyes, detergents, macrolides, aminoglycosides, and quinolones (Nishino et al., 2021).

Multiple studies have investigated the effect of acriflavine on *E. coli*, particularly in the context of antibiotic resistance, suggesting that the AcrAB-TolC pump actively expels it from the cell through an energy-dependent mechanism (Nakamura and Suganuma, 1972; D. Ma et al., 1993; Ma et al., 1995). Moreover, treatment with acriflavine also led to alterations in the structure of the plasma membrane (Nakamura, Yokomura and Hirayoshi, 1982; D Ma et al., 1993). *E. coli*, like many other Gram-negative bacteria, has intrinsic resistance to hydrophobic inhibitors such as acriflavine; while the outer membrane slows their influx, their equilibration time across the membrane is much shorter than the bacterial doubling time (Plésiat and Nikaido, 1992). This necessitates additional protective mechanisms, such as active efflux, to further defend against these compounds. Under stress conditions, like the presence of toxic compounds, the transcription of *acrAB* increases in *E. coli*, suggesting that a primary physiological function of the AcrAB complex is to protect the bacteria from various inhibitors, including acriflavine (Ma et al., 1995).

The susceptibility of phage T4 to acriflavine underscores the complex interplay between chemical agents and viral systems. While early studies provided foundational insights, contemporary research continues to refine our understanding of these interactions. Learning about phage susceptibility to chemical agents like acriflavine can offer insights into their role in phage-host interactions and phage resistance mechanisms and inform the development of better phage therapies.

In this work, we aim to investigate the role of the *ac* gene in phage T4 susceptibility to acriflavine by characterizing the effects of gene knockouts and modifications, and to elucidate the mechanism by which the Ac protein influences efflux pump function in *E. coli*. Furthermore, we seek to explore the evolutionary conservation of acridine resistance genes in phages from diverse species and their potential impact on phage-host interactions.

## Materials and Methods

### Bacteria and Bacteriophage Strains

Bacterial cultures of *Escherichia coli* CR63 (K strain, *supD,* F+, λ -*serU60*(AS), *lamB63)* provided by the Reha-Krantz lab at the University of Alberta, were used for phage amplification and plating. 5-alpha competent *Escherichia coli* from New England Biolabs was used for plasmid work and phage recombination. *Escherichia coli* Rosetta-gami B(DE3) (Novagen) (*F^-^ ompT hsdSB (rB^-^mB^-^)gal dcm lacY1 ahpC(DE3) gor522::Tn10 trxB pRARE (Cam^R^, Kan^R^, Tet^R^)* was used for plasmid expression and experiments. *Escherichia coli* bacteriophage T4 11303-B4 (ATCC 11303-B4) was used for this study. All bacterial cultures were grown on Luria-Bertani Miller broth (Fisher Scientific). Plating was done on HB agar (bacto tryptone 13g/L (Fisher Scientific), sodium chloride 8g/L (Fisher Scientific), sodium citrate 1g/L (BDH), glucose 1g/L (Fisher Scientific), bacto agar 6g/L (Fisher Scientific) for soft-agar and 12g/L for hard agar plates with and without acriflavine (1μg/ml) (Sigma-Aldrich).

### Determination of phage titers

A modified version of the soft agar overlay technique, detailed in Clokie and Kropinski, (2009), was used. Phage samples were serially diluted in Luria-Bertani Miller broth. 4 ml of molten soft agar was mixed with 0.1 ml of the overnight host culture and 0.1 ml of phage dilution. The mixture was poured over HB hard agar plates, with and without acriflavine, and incubated overnight at 37°C (incubator-shaker Ecotron, Infors HT). Individual phage plaques were counted, and titers were calculated as the average of plaque count number times the reciprocal of the dilution factor. Titers were reported in plaque-forming units per millilitre (PFU/ml).

### Ac vector

The Ac vector plasmid (**Figure S1**) was built from the pUC19 plasmid (Novagen) structure. Briefly, 200 bp of DNA homologous to the regions immediately up and downstream of the T4 *ac* gene were amplified by polymerase chain reaction (PCR) using Phusion High-Fidelity Polymerase (Thermo Fisher Scientific) in a T100 Thermal Cycler (Bio-Rad) from genomic phage T4 DNA using primer pairs MH69 and MH70, and MH71 and MH72 for the 5’ and 3’ homology regions, respectively (**Table S1, Table S2**). These DNA fragments were ligated into the HindIII and EcoRI sites of the pUC19 plasmid. A multiple cloning site with SacI, KpnI, BamHI, XbaI and SalI restriction enzyme sites was created between the 5’ and 3’ homology sequences by including these restriction sites in the PCR primers (**Table S1**).

### Modifications of phage TO ac gene

Deletion of the *ac* gene in phage T4 (T4Δ*ac*) was performed through homologous recombination with the Ac vector. In short, competent *E. coli* DH5α was used for cloning, plasmid propagation, and homologous recombination in Luria-Bertani Miller broth supplemented with 100 μg/ml of ampicillin. Homologous recombination was performed in triplicate. Bacterial host cultures were grown at 37 °C, with aeration until OD600 ∼0.2. Cells were then infected with phage T4 at a multiplicity of infection (MOI) of 0.01. Infections were carried out for 4 h at 37 °C, 250 rpm. When lysis was observed, phage lysates were filtered through a 0.22-μm syringe filter (Corning) and plated on HB and HB-acriflavine (HB-Acr) agar plates, as previously described. Acriflavine-resistant plaques were picked on HB-Acr selection plates and used to inoculate 2 ml of exponentially growing *E. coli* CR63 (OD600 ∼0.2). Infected *E. coli* CR63 cultures were grown at 37°C with aeration until complete lysis was observed. The resulting phage lysates were used directly in PCR to screen the *ac* gene region using primers LRK476 and LRK477 (**Table S1**, **Table S2**). PCR products were sent for Sanger DNA sequencing to confirm gene knockout.

A similar approach was used to find Tq phages carrying mutations (single-site insertion— Tqins, deletion—Tqdel, and substitution—Tqsub) in the *ac* gene. In this strategy, acriflavine-resistant plaques were picked on HB-Acr selection plates, amplified and validated in the same manner as for complete gene deletion described above.

### Expression of ac gene in E. coli

The wild-type *ac* gene sequence from phage T4 was amplified by PCR using primers SR06 and SR07 (**Table S1**, **Table S2**) from a phage lysate and cloned into pET11a using the *NdeI* and *BamHI* restriction sites to create the pET11a-*ac* vector. This vector was then recovered and transformed into the expression strain *E. coli* Rosetta-gami B(DE3). This strain was grown overnight in Luria-Bertani Miller broth supplemented with 100 μg/ml ampicillin at 37°C, with aeration. 1% culture from the overnight culture was grown to OD600 ∼0.5. The cell culture was induced with 0.1mM IPTG at 37°C overnight with aeration.

The effect of the *ac* gene on *E. coli* Rosetta-gami B(DE3) was tested as follows. Overnight cultures—not carrying the pET11a-*ac* vector (empty), carrying the vector but not induced (non-induced), and carrying the vector and induced (induced)—were serially diluted and plated on HB and HB-Acr plates and incubated overnight at 37°C. Colonies were counted for each culture under different plating conditions. Cell concentration was reported as colony forming units per millilitre (CFU/ml).

The effect of the ac gene expression in *E. coli* Rosetta-gami B(DEs) on phage infection was tested as follows. Phage lysates of Tq and TqΔ*ac* infections were plated on HB and HB-Acr plated. The lysates were serially diluted, and t.u ml were mixed with molten soft agar and t.u ml of overnight cultures of *E. coli* Rosetta-gami B(DEs)—empty, non-induced, and induced. Plates were incubated overnight at sv°C. Plaques were counted in the presence and absence of acriflavine with the different strains as bacterial hosts. Titers were reported in plaque forming units per millilitre (PFU/ml).

### Predicted protein structure

The predicted structure of Ac, the *ac* gene product, was obtained using ESMfold (Lin *et al*., 2023).

### BLAST Analysis

Basic Local Alignment Search Tool (BLAST) *(Altschul* et al.*, UVVW)* analysis (https://blast.ncbi.nlm.nih.gov/Blast.cgi) was conducted to identify putative homologous sequences of the *ac* gene from phage Tq in the NCBI GenBank database *(Zhang* et al.*, YWWW; Morgulis* et al.*, YWW[)*. BLASTN x.uy.u+ (for the nucleotide sequence) was performed using default parameters, with the phage *ac* gene sequenced from this study as the query against the GenBank non-redundant database. To improve confidence in homology inference, BLASTN hits were evaluated holistically, considering bit score, *E*-value, percent identity, and alignment coverage together, rather than relying on any single metric. *E*-value served as the primary criterion, with hits in the range of ut^-0^ to ut^-12^ retained as putative homologs (Pearson, xtus). A high query coverage threshold of zt% (Janaki, Gowri and Srinivasan, xtxu) was considered to ensure substantial alignment *(Nestor* et al.*, YWY\)*. As such, hits with low query coverage or unusually low percent identity despite a significant *E*-value were manually examined to prevent false positives. The results were filtered to retrieve the top hits corresponding to phages that infect different bacterial genera to ensure a broader representation of phage diversity across various hosts. Multiple sequence alignments (MSA) were generated using MUSCLE vs.z.su (Edgar, xttq) with default parameters. Conserved regions were visualized using Jalview x.uu.q.u.

## Results

To understand the potential mechanism through which acriflavine affects the phage T4 infection process, different conditions were tested. **Figure 1** shows a graphic representation of the bacterial strains and phage variants used in this study.

**Figure 1.**
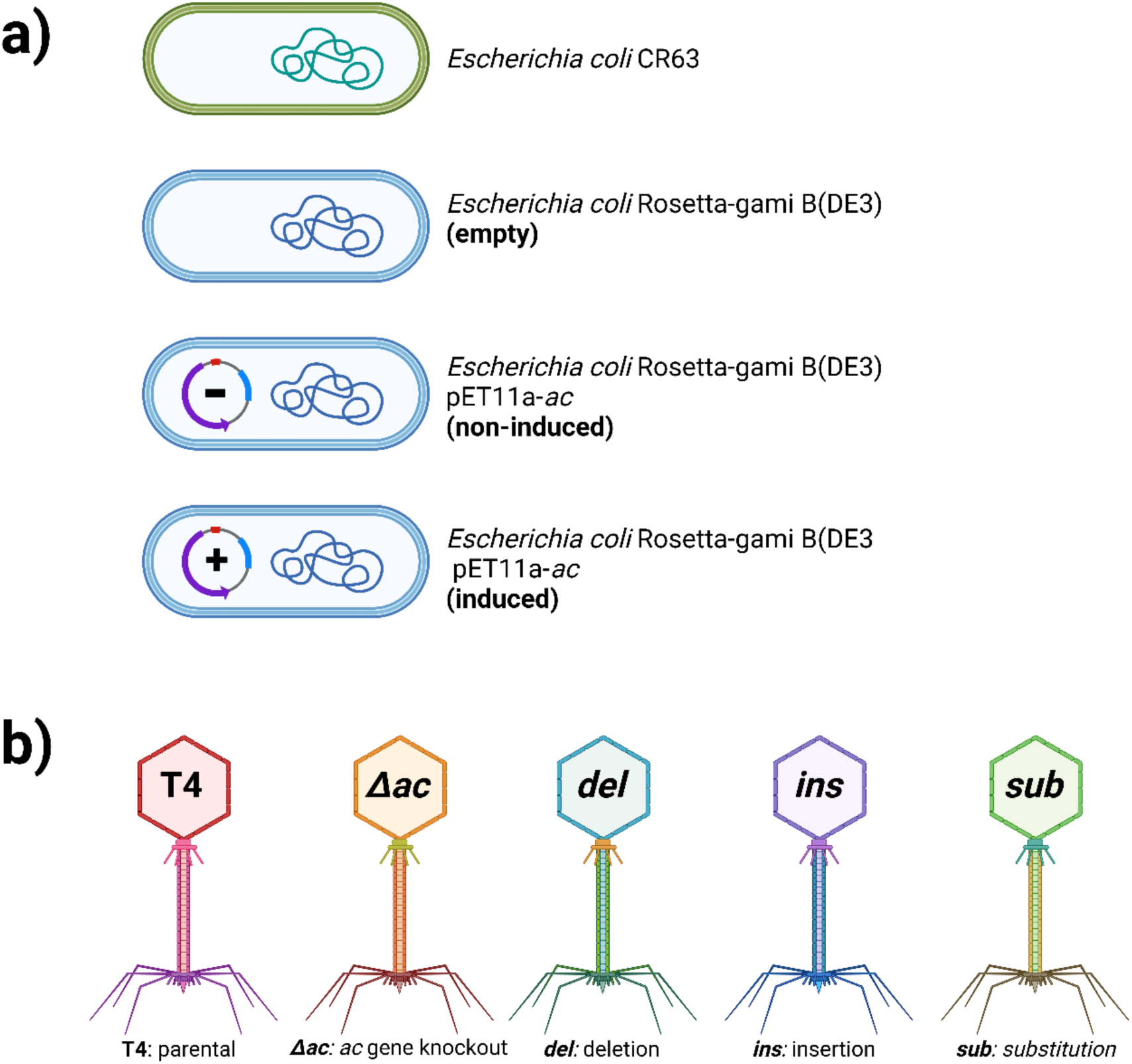
a) Bacterial strains used in this study: *E. coli* CR63 for amplifying and plating T4 phage variants. *E. coli* Rosetta-gami B(DE3) (empty), *E. coli* Rosetta-gami B(DE3) carrying the plasmid pET11a-ac (non-induced), and *E. coli* Rosetta-gami B(DE3) carrying the plasmid pET11a-ac induced with IPTG.(induced) b) Phage variants used: Parental T4 and T4 with the knockout of the ac gene (T4Δac), T4 carrying an insertion in the ac gene sequence (T4_ins_), T4 carrying a deletion in the ac gene sequence (T4_del_), and T4 carrying a substitution in the ac gene sequence (T4_sub_). Created in BioRender. Arvizu, A. (2024) <u>BioRender.com/j35d082</u>

### Effect of acriflavine on TO and TO variants

Plaque counts of stocks of the parental phage T4 and the different phage variants (T4Δ*ac*, T4del, T4ins, T4sub) at the same titer were performed on HB plates containing acriflavine (**Figure 2**; **Table 1**). On the other hand, all phage lysates of purified variants formed plaques at ∼10^10^ PFU/ml titer on HB-Acr agar plates, four orders of magnitude greater than the titer observed with the parental T4 phage lysate on the same plates (2.39 x10^6^ PFU/ml). This low level of resistance in the parental phage has been observed previously (Hessler, 1963; Hessler, Baylor and Baird, 1967), and corresponds to background mutations naturally occurring in the population.

**Figure 2.**
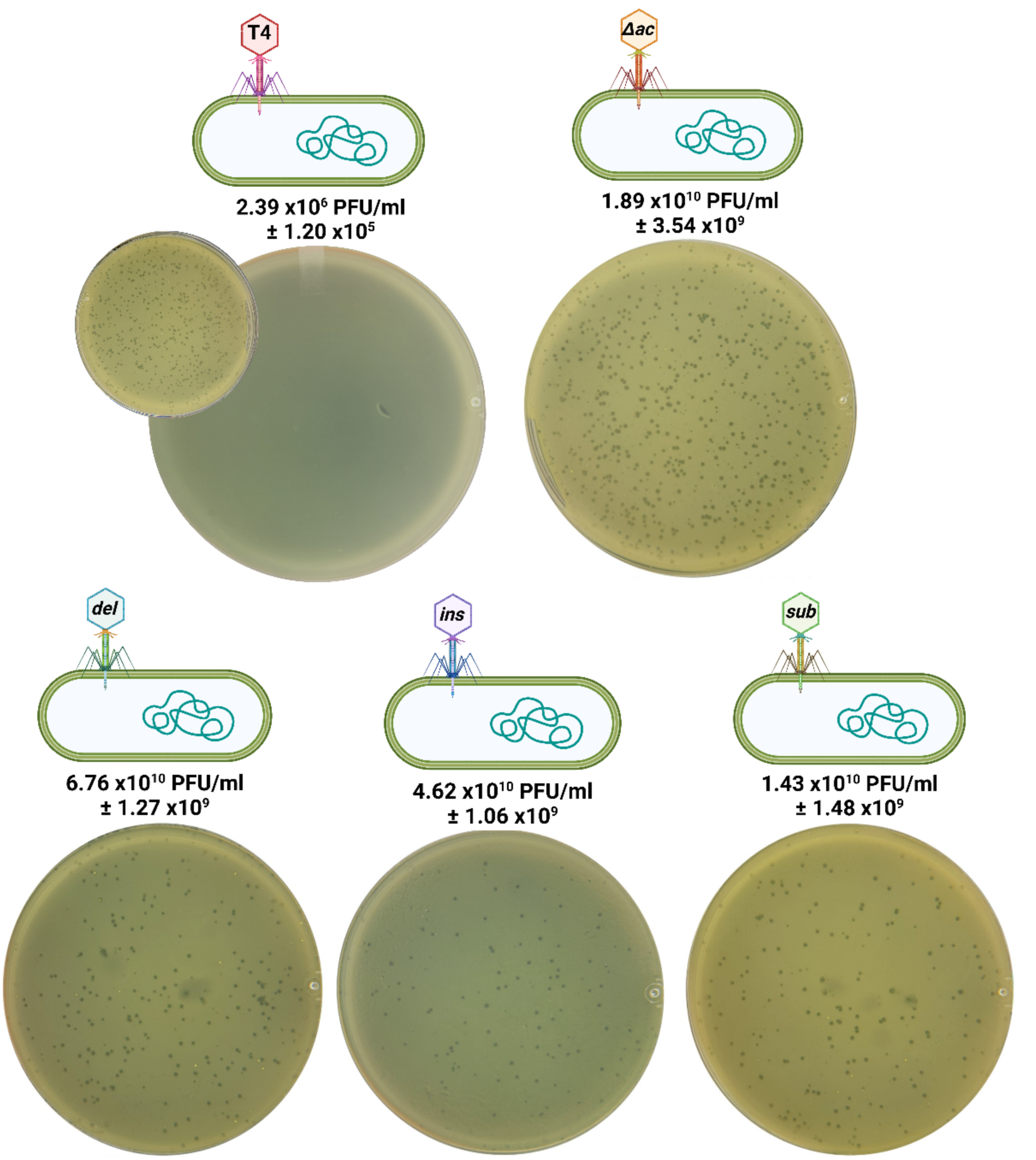
Titers of phage T4 variants—T4, T4Δac, T4_del_, T4_ins_, T4_sub_ —plated on HB with acriflavine using *E. coli* CR63 as host. Plates shown are for 10^8^ dilutions of phage lysate. Inset plate for parental T4 shows 10^4^ dilution of the phage lysate. Created in BioRender. Arvizu, A. (2024) <u>BioRender.com/j01g065</u>

**Table 1.** Table 1 Titers of T4 phage variants on HB-Acr plates from results shown in Figure 2.

| Phage Variant | DNA sequence change | Amino acid change (three-letter code) | Type of mutation | Titer on HB-Acr*<br>(PFU/ml) |
| --- | --- | --- | --- | --- |
| T4 | - | - | - | $2.39 \times 10^6$<br>$\pm 1.20 \times 10^5$ |
| T4Δac | Δac | - | Knockout | $1.89 \times 10^{10}$<br>$\pm 3.54 \times 10^8$ |
| T4<sub>del</sub> | c.38_39delT | p.Cys14ValfsX2 | Frameshift | $6.76 \times 10^{10}$<br>$\pm 1.27 \times 10^9$ |
| T4<sub>ins</sub> | c.47_48insA | p.Val17SerfsX12 | Frameshift | $4.62 \times 10^{10}$<br>$\pm 1.06 \times 10^9$ |
| T4<sub>sub</sub> | c.112G>C | p.Gly38Arg | Missense | $1.43 \times 10^{10}$<br>$\pm 1.48 \times 10^9$ |

\**E. coli* CR63 was used as the bacterial host.

### Effect of acriflavine on E. coli variants with and without the ac gene

The susceptibility of *E. coli* Rosetta-gami B(DE3) and its variants—empty, non-induced, and induced—to acriflavine was assessed by plating culture dilutions on HB agar plates with and without acriflavine (**Figure 3**). In the absence of acriflavine, the colony counts were 2.28 x10^9^ CFU/ml for the empty strain culture, 4.33 x10^9^ CFU/ml for the non-induced culture, and 1.51 x10^9^ CFU/ml for the induced culture. However, in the presence of acriflavine, the induced culture showed a decrease in growth of three orders of magnitude (down to 1.41 x10^6^ CFU/ml), while the other two cultures—empty and non-induced—showed a reduction of less than one log (to 6.7 x10^8^ CFU/ml and 4.72 x10^8^ CFU/ml, respectively).

**Figure 3.**
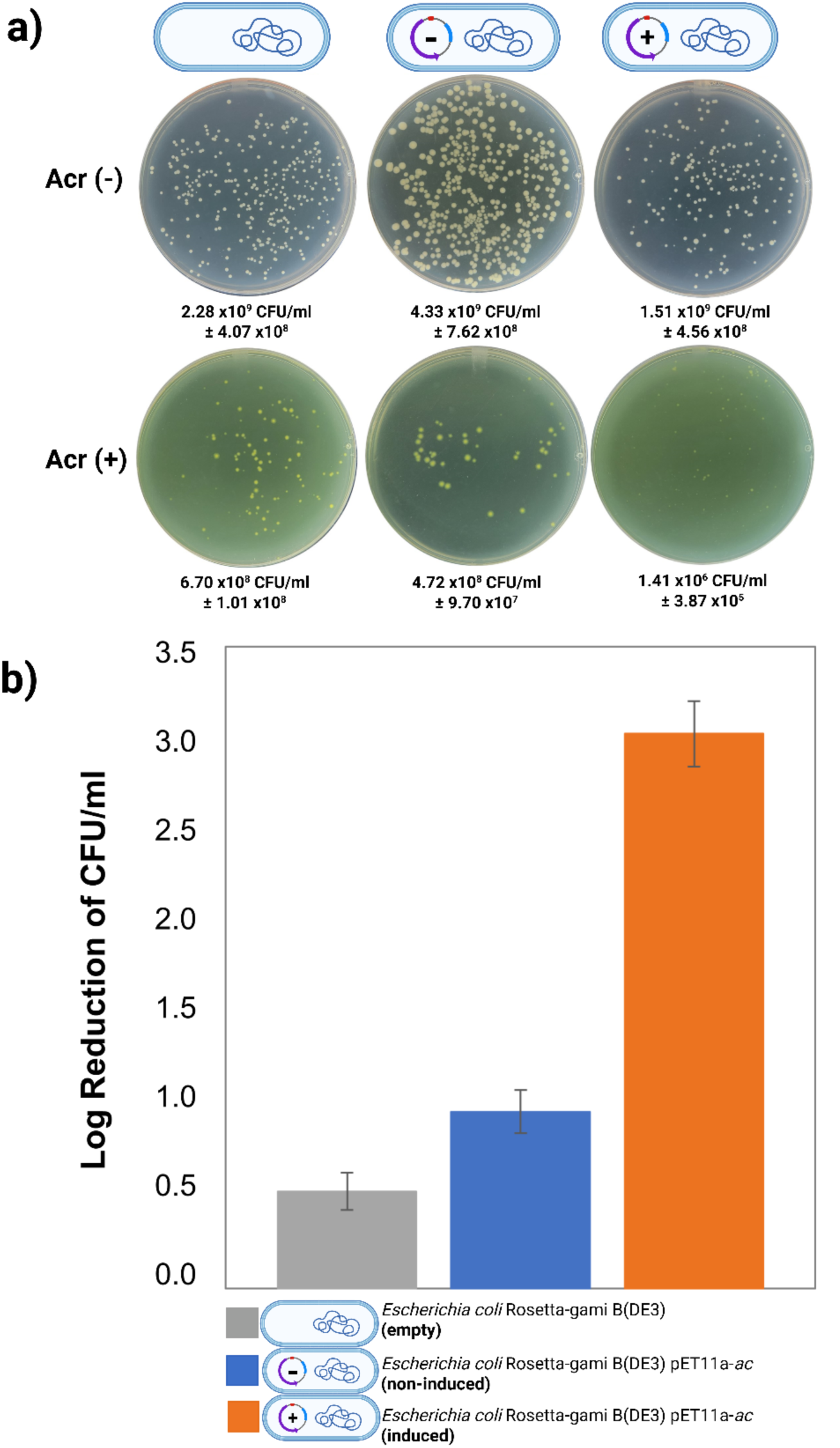
Effect of acriflavine on the growth of *E. coli* Rosetta-gami B(DE3) variants. a) Plates of HB agar showing the colonies for dilution 10^7^ without (Acr(-))and with (Acr(+)) acriflavine for dilution 10^7^ of the strain not carrying the plasmid (empty), carrying the plasmid but not expressing the gene (non-induced), and dilution 10^4^ for strain carrying the plasmid and expressing the *ac* gene (induced). b) Log reduction of CFU/ml of the different bacterial strain variants in the presence of acriflavine. Created in BioRender. Arvizu, A. (2024) <u>BioRender.com/y59q690</u>

### TO phage infection of E. coli variants

Infections of the variants of *E. coli* Rosetta-gami B(DE3)—empty, non-induced, and induced—were conducted on HB and HB-Acr plates with phages T4 and T4Δ*ac* in order to assess the impact of the Ac protein (**Figure 4**).

**Figure 4.**
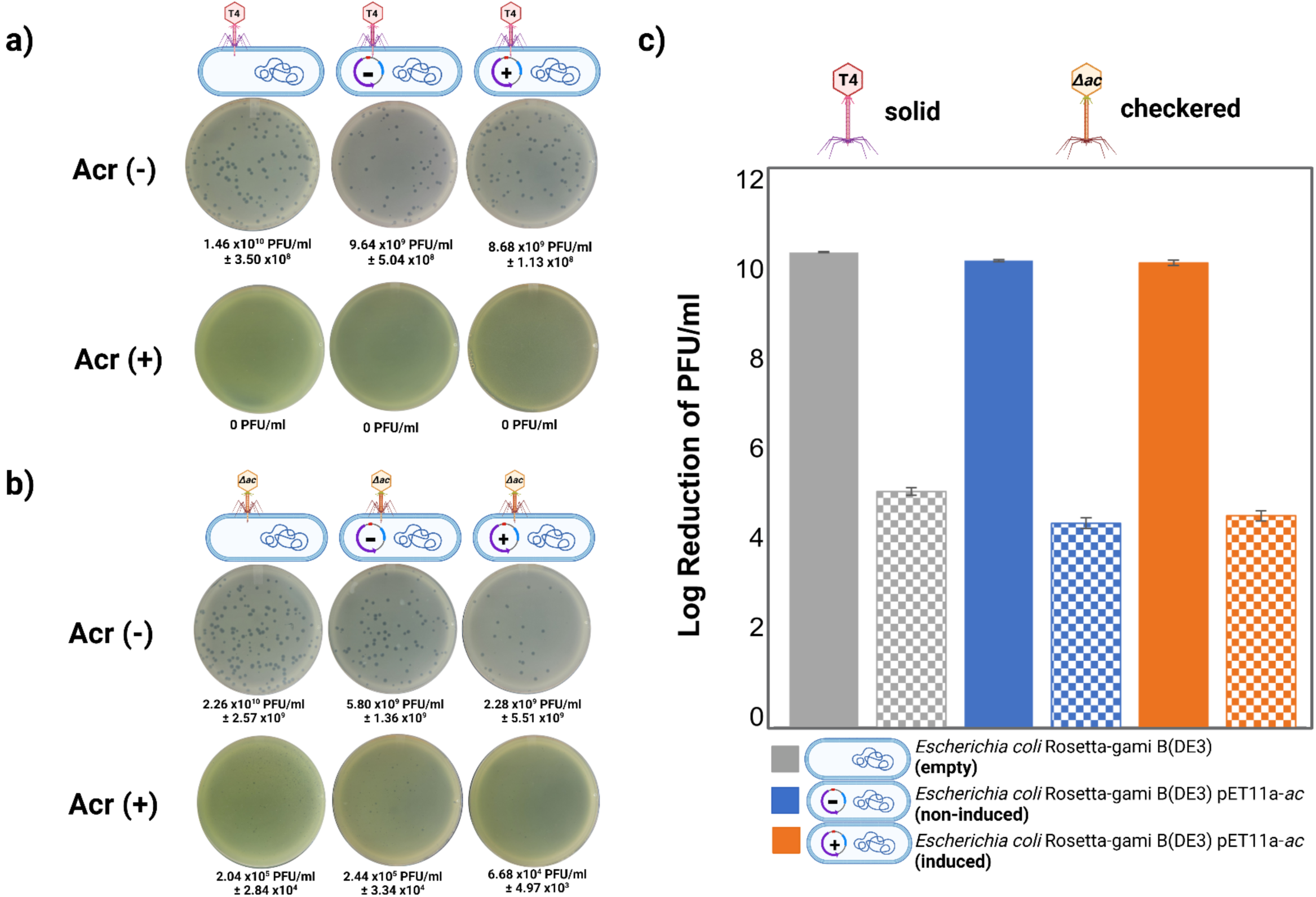
Phage titers with different variants of *E. coli* Rosetta-gami B(DE3) as hosts. a) Plates showing plaques for dilution 10^8^ of phage lysates on HB agar without (Acr(-)) and with (Acr(+)) acriflavine for dilution 10^2^ of parental T4 b) Plates showing plaques for dilution 10^8^ on HB agar without (Acr(-)) and with (Acr(+)) acriflavine for dilution 10^3^ of T4Δac c) Log reduction of PFU/ml in the presence of acriflavine of both phage variants with the different strain variants as hosts. Created in BioRender. Arvizu, A. (2024) <u>BioRender.com/x60w878</u>

In the case of the parental T4, the titer in the absence of acriflavine was 1.46 x10^10^ PFU/ml when infecting the empty host, 9.64 x10^9^ PFU/ml when infecting the non-induced host, and 8.68 x10^9^ PFU/ml when infecting the induced host. However, in the presence of acriflavine, T4 phage was not able to form plaques in either of the three variant hosts. In contrast, while the titers of T4Δ*ac* phage infections in the absence of acriflavine were 2.26 x10^10^ PFU/ml for the empty host, 5.8 x10^9^ PFU/ml for the non-induced host and 2.28 x10^9^ PFU/ml for the induced host, the phages retained activity when infecting in the presence of acriflavine, with titers of 2.04 x10^5^ PFU/ml, 2.44 x10^5^ PFU/ml, and 6.68 x10^5^ PFU/ml, for each host respectively. One-way ANOVA showed no significant difference (p > 0.05) in titers between the two phage variants—parental T4 and T4Δ*ac*— across the three hosts variants in the absence of acriflavine.

### BLAST Analysis

BLASTN analysis of the *ac* gene sequence against the full NCBI database identified 289 hits, with sequence identities between 93 to 100%. These hits correspond to phages infecting host from diverse bacterial genera, including *Escherichia*, *Yersinia*, *Salmonella*, *Shigella*, *Pseudomonas, Citrobacter*, and *Serratia*. The Multiple Sequence Alignment (MSA) of the DNA sequences (**Figure S3**) and amino acid alignments (**Figure 5a**) generated using MUSCLE highlights the high conservation of the *ac* gene and Ac protein across phages targeting this broad range of hosts. The analysis also includes the T4 variants engineered in this study, phage T2 and phage PP01 from our lab stocks. Furthermore, **Figure 5b** presents the predicted 3D structure of the *ac* gene product, modeled using ESMfold.

**Figure 5.**
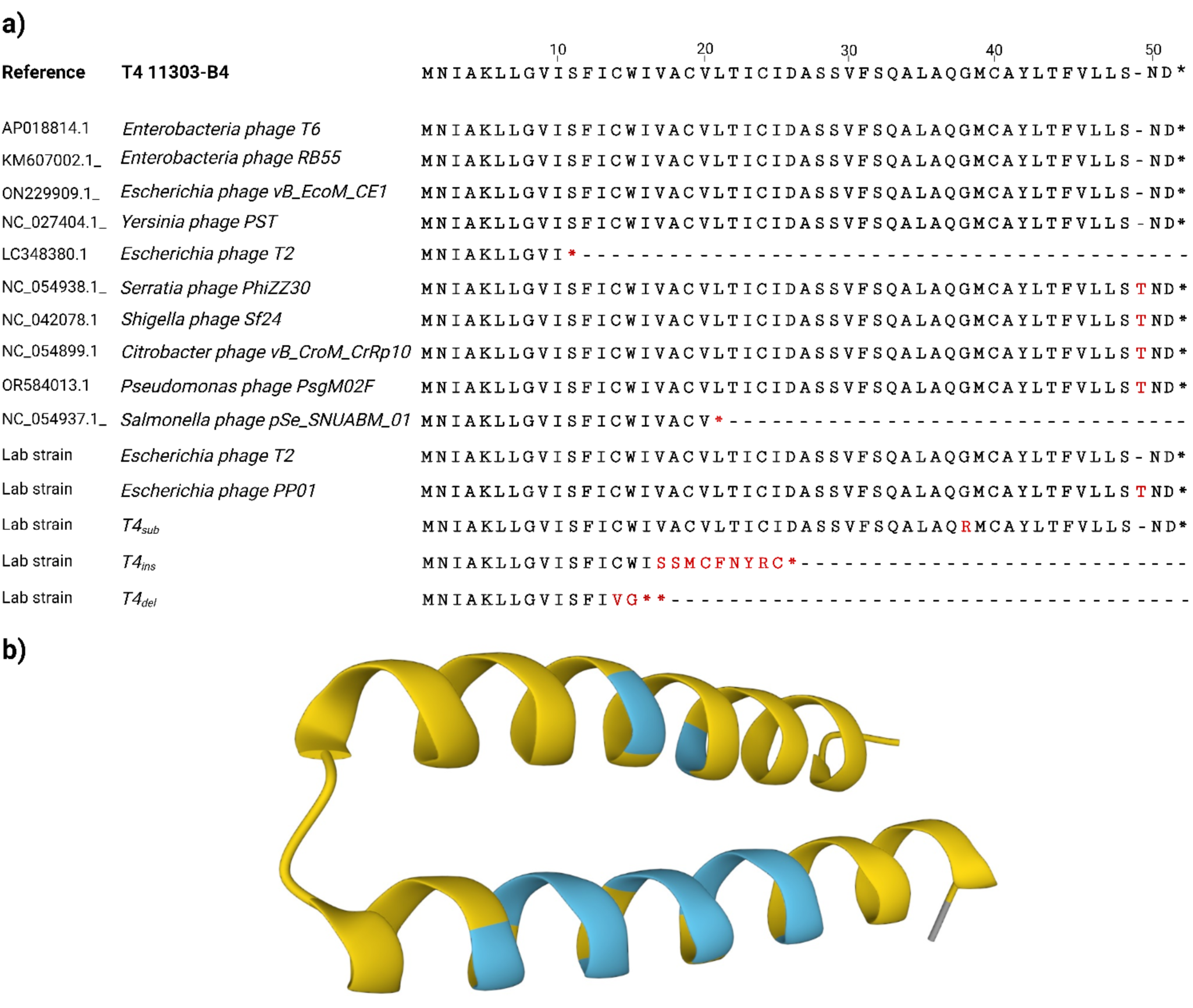
Conservation of Ac, the *ac* gene product, among various bacteriophages. a) Multiple Sequence Alignment (MSA) by MUSCLE of the amino acid sequences of the *ac* gene product across diverse phages. It should be noted that phage T2 is represented twice: LC348380.1 is the reference sequence provided by NCBI, while the Lab strain sequence was obtained experimentally from a phage T2 cultured in our lab. The * symbol designates a stop codon. b) Predicted protein structure of the *ac* gene product Ac generated using ESMfold. Created in BioRender. Arvizu, A. (2025) https://BioRender.com/p31z519

## Discussion

The comparison of infections of *E. coli* CR63 by the parental phage T4 and the T4 *ac* variants—whether knockout, deletion, insertion, or substitution—in the presence of acriflavine clearly demonstrated the impact of the *ac* gene on acriflavine susceptibility (**Figure 2**). These results align with previous studies suggesting that acridine play a significant role in infections of bacteria by phages T2, T4, and T6 (Fitzgerald and Lee, 1946; Foster, 1948; Silver, 1964; Hessler, 1965); in these studies, the authors proposed that infected hosts absorbed more acridine while bacteria infected by acriflavine-resistant phages exhibited reduced uptake, allowing phages to replicate and produce new progeny. Our work proposes a different mechanism is at play, in which Ac does not promote the uptake of acridines but rather interacts with efflux pumps to impede their removal from the cells (more details provided below). These early studies also showed differential susceptibility between the T-even phages to proflavine, an acriflavine derivative, with T6 being more sensitive than T2 and T4 (Foster, 1948). This difference in susceptibility is thought to be related to variations in the phage DNA or differences in the response of the host cell to different acridines (Silver, 1964; Tubbs, Ditmars and Van Winkle, 1964). It should also be noted that acriflavine has been shown to specifically inhibit the replication of certain plasmids *in vivo*, without affecting that of the *Escherichia coli* K-12 chromosome (Nakamura, 1965, 1974, 1976). In addition to further confirming the connection between the *ac* gene and acriflavine resistance, the results in the present study demonstrate how point mutations or knockouts affect this resistance and influence phage infections.

Expression of the *ac* gene from a plasmid transformed into *E. coli* Rosetta-gami B(DE3) enabled the investigation of the effect of the *ac* gene product, Ac, itself on the bacterium in the presence of acriflavine (**Figure 3**). While cultures of the parental strain (empty) and of the non-induced strain each showed a 0.5-log and a 1-log reduction in CFU/ml counts, respectively, in the presence of acriflavine, this effect was substantially greater (3-log reduction) in the cultures expressing the *ac* gene product (**Figure 3b**). While acriflavine is known to inhibit bacterial growth, the greater reduction in CFU/ml when the *ac* gene is expressed may be due to the Ac protein altering the function of the AcrAB-TolC efflux pump, reducing its efficiency in extruding acridines from the cell. This hypothesis aligns with findings by Silver (1967), which suggest that the *ac* gene (previously referred to as *pr* gene) product from phage T2—sharing over 96% sequence identity with the *ac* gene from T4—modifies the membrane permeability by either integrating itself into the membrane or interacting with existing membrane components, which include the AcrAB-TolC efflux pump. The predicted structure of the Ac protein (ESMfold prediction shown in **Figure 5b**), with two short alpha helices connected by the amino acids aspartate (D) and alanine (A) at positions 26 and 27, respectively, suggests it is membrane-associated. While this is consistent with a potential increased uptake of acriflavine, our results suggest it is more likely to interact with EP systems, hindering their function. In fact, host cells expressing the *ac* gene (induced) exhibited a significant sensitivity to ampicillin (100 μg/ml) on HB agar plates, both in the presence and absence of acriflavine (**Figure S2**). After 20 h of incubation at 37°C, the strain expressing the Ac protein exposed to ampicillin showed a 4-log decrease in growth compared to cells harbouring the plasmid but not expressing the *ac* gene (non-induced) under the same conditions. In *E. coli*, ampicillin is primarily hydrolysed by the β-lactamases encoded by genes such as *blaTEM, blaSHV, blaOXA*, among others. Additionally, the AcrAB-TolC EP contributes to the overall resistance mechanism by extruding the antibiotic, reducing its intracellular concentration, and potentially decreasing its effectiveness (Singh *et al*., 2019; Smith, Fernando and King, 2024). These findings suggest that the *ac* gene product possibly modifies or obstructs efflux pumps, thereby leading to an accumulation of antibiotic (and acridines) in the cell and increasing its susceptibility to these compounds.

This being said, even if the AcrAB-TolC pump is impaired due to the presence of the Ac protein, *E. coli*, like many other bacteria, possesses multiple systems to ensure survival under various environmental conditions, including exposure to inhibitors such as acriflavine. One class of these systems is the ATP-Binding Cassette (ABC) transporters, which use the energy from ATP hydrolysis to export drugs and various other molecules (Davidson *et al*., 2008). In a study by (Lee *et al*., 2003), the EfrAB pump, an ABC transporter from *Enterococcus faecalis*, was cloned into the drug-hypersensitive *E. coli* strain KMA32, which is deficient in the AcrAB pump. The transformed strain exhibited increased resistance to acriflavine, indicating that AcrAB-TolC is not the sole defence mechanism in *E. coli* against compounds like acriflavine (Lee *et al*., 2003; Teelucksingh, Thompson and Cox, 2020).

The different patterns of infection with different *E. coli* strain variants also provide insights on the Ac mechanism of action. The findings of Nakamura and Suganuma, (1972), who showed acriflavine-induced modifications of *E. coli* cell surface composition or structure, could explain the reduced infection by phages in the presence of the acridine; for example by such modifications potentially reducing the availability of phage receptors. The long tail fibre of the T4 phage binds to the outer membrane of *E. coli* using lipopolysaccharides (LPS) and OmpC as receptors (Washizaki, Yonesaki and Otsuka, 2016; Islam *et al*., 2019). Conversely, *E. coli* strain B has a deletion in the *ompC* sequence which causes the T4 phage to rely solely on LPS as a receptor when infecting this strain (Wilson *et al*., 1970; Prehm *et al*., 1976; Montag *et al*., 1990). As shown in **Figure 4**, in the absence of acriflavine, phage T4 can infect the B-type strain, *E. coli* Rosetta-gami B(DE3) — whether empty, non-induced, or induced — yielding similar viral titers. However, addition of acriflavine prevented the infection. This is likely due to an increase in intracellular acriflavine concentration inhibiting phage replication (Foster, 1948; Silver, 1964) (as described above) and, possibly, the potential reduction of available phage receptors on the cell surface as proposed by (Nakamura and Suganuma, 1972). However, in our experiments, T4Δ*ac*, which has the same receptor-binding proteins as T4, was able to infect the same host in the presence of acriflavine whether the host was the empty, non-induced or induced variant, as seen by the plaques obtained on HB-Acr agar plates (**Figure 4b**). In this case, the fact that the T4Δ*ac* was still able to infect and form plaques on the induced *E. coli* host, expressing Ac, in the presence of acriflavine strongly suggests that cell surface modification and reduced receptor availability is not the main mechanism impeding phage replication. However, it appears that the Ac protein is directly involved in making the phage more vulnerable, as observed by the reduced titer when compared to the other host variants (**Figure 4b**). This may indicate that while the Ac protein interaction is still occurring, it is less efficient when produced by the host rather than the phage, possibly due to differences in expression timing, or the protein localization or folding. Notably, viral titers on HB agar (acriflavine-free) plates were not affected regardless of the strain used as the host.

This being said, the lack of OmpC in the B strain has an impact on the phage’s ability to infect the host effectively in the presence of acriflavine. This is evidenced by the fact that the same phage—(T4Δ*ac*)—can form plaques in the 10^10^ PFU/ml range when infecting *E. coli* strain CR63, a K-12 strain, under the same experimental conditions, and the background of naturally occurring mutations of the T4 phage lysate can attach and infect this strain too, yielding a viral titer of 2.36 x10^6^ PFU/ml (**Figure 2**).

Finally, it is important to note that the *ac* gene is not exclusively found in phage T4. In fact, many other phages possess gene homologs of the T4 *ac* gene (**Figure 5, Figure S3**), suggesting a conserved function of the gene products based on amino acid identity. The small differences in identity between these genes in different phages may reflect evolutionary adaptations to specific bacterial hosts or environmental pressures; however, the fact that single-point mutations within the gene can lead to resistance to acridines also suggests high specificity. In fact, the presence of homologs across multiple phage species infecting bacteria of different genera and the high level of sequence conservation —particularly in key regions of the gene products— highlight the evolutionary significance of this gene. Despite the relatively short sequence, synonymous point mutations that do not alter the encoded amino acid sequence are prevalent. Additionally, a common codon insertion is observed three codons upstream of the stop codon resulting in the addition of a threonine (T) residue to the protein sequence in some phages. Furthermore, our lab stock of phage PP01, which carries the additional threonine residue, exhibited no changes in susceptibility to acriflavine, suggesting that the insertion does not confer resistance to the acridine compound. In contrast, the T4sub variant, which carries a point mutation leading from a glycine (G) to arginine (R) substitution at residue 38, demonstrates resistance to acriflavine, emphasizing the functional impact of specific amino acid changes on the Ac function.

In this context, the fact that the *ac* gene confers susceptibility, rather than resistance, to an active agent is, in itself, an interesting evolutionary contrast. One hypothesis is that this mechanism would play a role in reducing the rate of infection within a population in which high levels of lysis occur, which leads to the release of natural molecules with structures or functions similar to acridines. These include phenazines, which can act as electron shuttles, contribute to biofilm formation, act as signals for gene expression, and increase bacterial virulence (Pierson and Pierson, 2010), and quinones, which are active participants in the electron transfer chain (van Beilen and Hellingwerf, 2016). In this case, the release of these compounds through cell lysis would lead to halted or tempered infection of the strictly lytic phage within the host population to enable longer-term propagation of the phage (avoiding rapid host extinction). This would have similarities to phage quorum sensing strategies previously described (León-Félix and Villicaña, 2021). In this context, the Ac protein would act as a control/response mechanism to thwart the rapid progression of phage infection under specific conditions. This provides a foundation for further functional studies to explore the role of acridine resistance genes in phage biology, including their contribution to phage fitness, infection efficiency, and their influence on host resistance mechanisms.

## Conclusion

This study provides valuable information on the nature and mechanism of action of the Ac protein found in multiple phages. The work showed that knocking out or introducing single mutations in the *ac* gene of phage T4 makes the mutant phages resistant to acriflavine. When the *ac* gene was expressed in *E. coli*, our results showed that the Ac protein likely integrates into the bacterial membrane, potentially interacting with the AcrAB-TolC efflux pump, disrupting its function in expelling the acridine from the cell. Additionally, phage infections were altered in the presence of the *ac* gene product: parental phage T4 was unable to infect *E. coli* expressing the Ac protein in the presence of acriflavine, whereas the knockout mutant (T4Δac) successfully formed plaques under the same experimental conditions. Moreover, BLASTN analysis revealed high sequence conservation of acridine resistance genes across phages targeting various bacterial hosts, highlighting their evolutionary importance and significant role in phage-host systems. Identifying the Ac protein as a key factor in phage susceptibility opens the door to discovering new phage-encoded regulators of bacterial defenses and stress responses. The widespread conservation of *ac* across diverse phages suggests it may serve a broader function beyond acriflavine susceptibility, potentially influencing bacterial metabolism through efflux pump modulation. Further investigation into *ac* and related genes will deepen our understanding of how phages manipulate host defenses, with implications in microbial ecology, antibiotic resistance, and phage evolution.

## Acknowledgements

Funding for this work has been provided by the NSERC Discovery program, the Canada Foundation for Innovation John R. Evans Leaders Fund, the Alberta Innovates Accelerating Innovations into CarE (AICE) Concepts program, the Alberta Agriculture and Forestry Research and Development program, and CONACyT.

## Notes

### Competing Interest Statement

The authors have declared no competing interest.

