## Supplemental Materials for "The role and effects of the phage T4 Ac protein on infection"

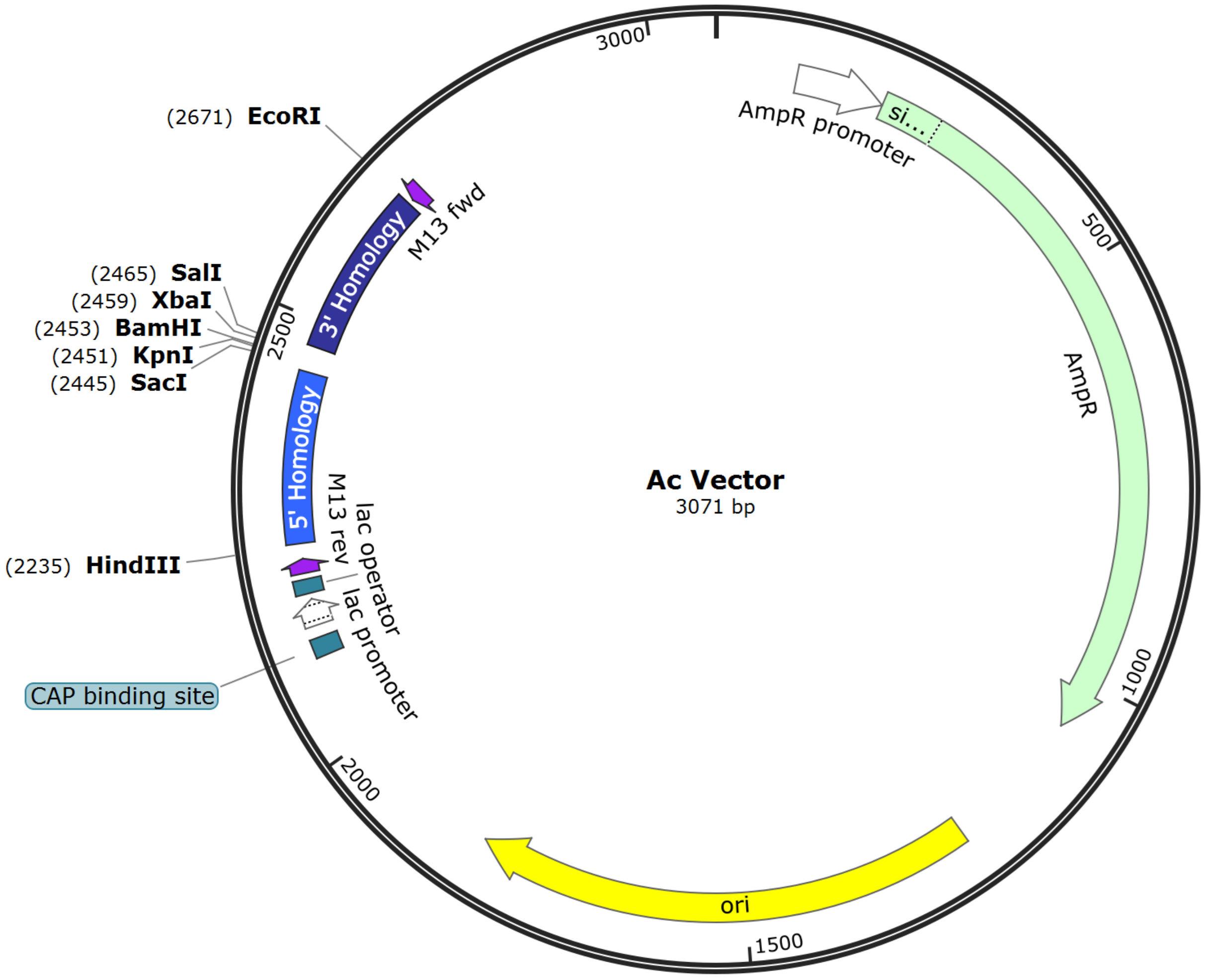


**Figure S1** Ac vector map.

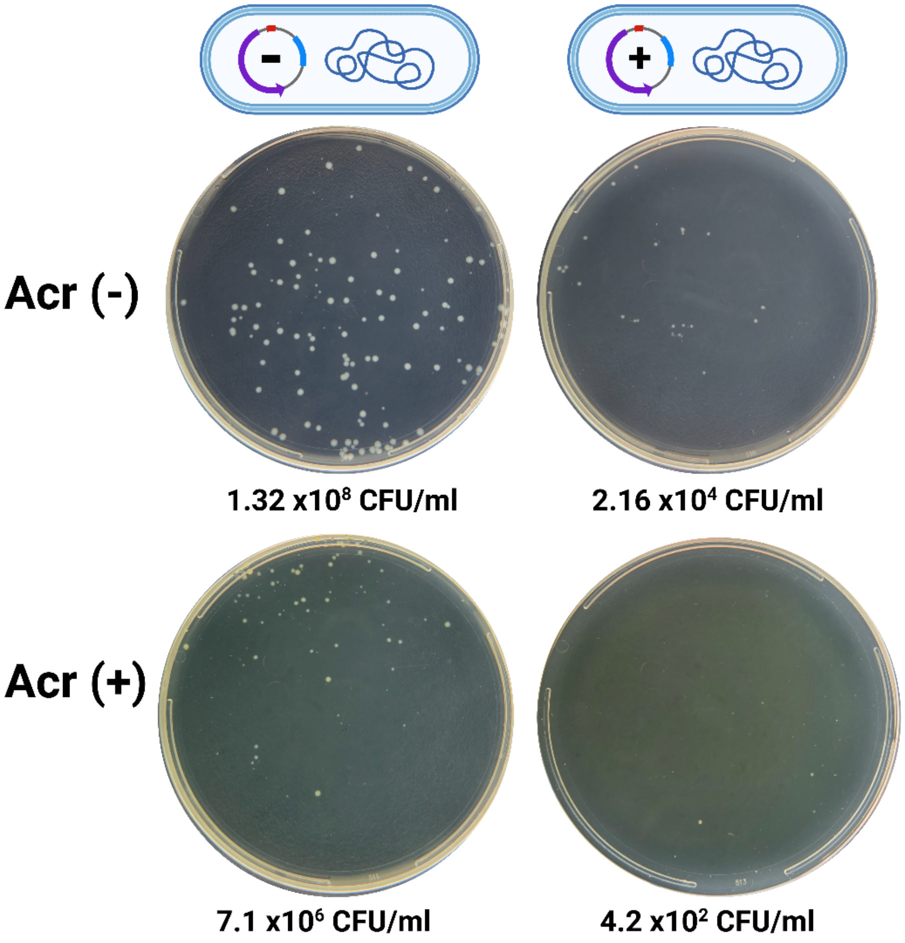


**Figure S2** Effect of ampicillin on Escherichia coli Rosetta-gami B(DE3) variants. Plates of HB-ampicillin agar without acriflavine (Acr(-)) showing colonies for dilution $10^5$ and with acriflavine (Acr(+)) for dilution $10^6$ of the strain not expressing the plasmid (non-induced) and for dilution $10^3$ (Acr(-)) and $10^2$ (Acr(+)) of the strain expressing the Ac protein (induced). Created in BioRender. Arvizu, A. (2024) BioRender.com/y77l729

**Table S1** Primer sequences

| Primer | Sequence 5' to 3' |
|---|---|
| MH69 | GCACCGTAGCAAGCTTAAGGTCAGCTAGCTTTGGCGTTTAC |
| MH70 | CAGGTTCGGATCCGGTACCGAGCTCATTTTCCTCACTGGCGTCCGAA |
| MH71 | GCACCGTAGCGGATCCTCTAGAGTCGACATGATTAAGAAAATCTTGGGCT |
| MH72 | CAGGTTCGAATTCAAGCCAATGCTTCATTATCAAT |
| LRK476 | CTACCAATAAAGCAGCAAGGGC |
| LRK477 | CCTCTAACGACGGGATTTGG |
| SR6 | CATATGATGAATATTGCAAAATTATTAGGAGT |
| SR7 | GGATCCTTAATGGTGATGGTGATGGTGG |

**Table S2** PCR conditions

| Step | Temperature | Time | Cycles |
|---|---|---|---|
| Initial Denaturation | 98 °C | 2 min | 1 |
| Denaturation | 98 °C | 30 sec | 35 |
| Annealing | 60 to 66 °C dependent on the primer set | 30 sec | |
| Extension | 72 °C | 15 sec/kb dependent on the amplicon length | |
| Final Extension | 72 °C | 10 min | 1 |

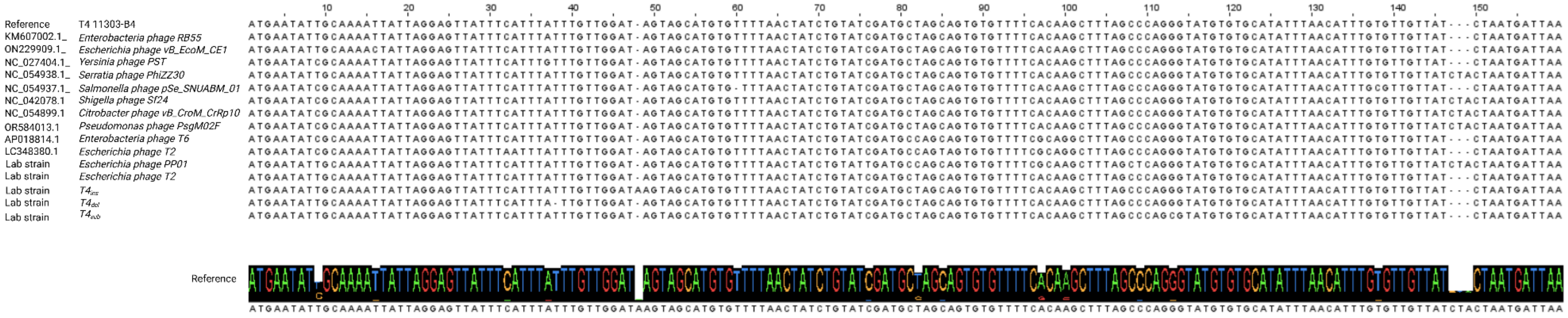


**Figure S0** Multiple sequence alignment (MSA) of the *ac* gene using MUSCLE. MSA shows gene conservation of the top hits across bacterial host genera from BLASTN, as well as T4 variants—$T4_{ins}$, $T4_{del}$, $T4_{sub}$— and phage T2 and phage PP01 lab stocks.
Created in BioRender. Arvizu, A. (2025) https://BioRender.com/p31z519
